# A highly expressed odorant receptor from the yellow fever mosquito, AaegOR11, responds to (+)- and (-)-fenchone and a phenolic repellent

**DOI:** 10.1101/2022.09.26.509539

**Authors:** WeiYu Lu, Walter S. Leal, Katherine K. Brisco, Sunny An, Anthony J. Cornel

**Author notes:** Corresponding author, (WSL).

## Abstract

The cornerstone of the reverse chemical ecology approach is the identification of odorant receptors (OR) sensitive to compounds in a large panel of odorants. In this approach, we de-orphanize ORs and, subsequently, measure behaviors elicited by these semiochemicals. After that, we evaluate behaviorally active compounds for applications in insect vector management. Intriguingly, multiple ORs encoded by genes highly expressed in mosquito antennae do not respond to any test odorant. One such case is CquiOR125 from the southern house mosquito, *Culex quinquefasciatus* Say. To better understand CquiOR125’s role in *Culex* mosquito olfaction, we have cloned a CquiOR125 orthologue in the genome of the yellow fever mosquito, *Aedes aegypti* (L.), AaegOR11. Unlike the unresponsive nature of the orthologue in *Cx. quinquefasciatus*, oocytes co-expressing AaegOR11 and AaegOrco elicited robust responses when challenged with fenchone, 2,3-dimethylphenol, 3,4-dimethylphenol, 4-methycyclohexanol, and acetophenone. AlphaFold models showed that AaegOR11 and CquiOR125 share structural homolog cores with MhraOR5, the only insect OR structure (PDB: 7LID) elucidated to date. Interestingly, AaegOR11 responded strongly and equally to (+)- and (-)-fenchone, with no chiral discrimination. Contrary to reports in the literature, fenchone did not show any repellency activity against *Ae. aegypti* or *Cx. quinquefasciatus*. Laboratory and field tests did not show significant increases in egg captures in cups filled with fenchone solutions compared to control cups. The second most potent ligand, 2,3-dimethylphenol, showed repellency activity stronger than that elicited by DEET at the same dose. We, therefore, concluded that AaegOR11 is a mosquito repellent sensor. It is feasible that CquiOR125 responds to repellents that remain elusive.

## Introduction

The explosive advancement in our understanding of the molecular basis of insect olfaction in the last two decades paved the way for the application of reverse chemical ecology [1]. In conventional chemical ecology approaches, attractants, repellents, and other semiochemicals are chemically characterized after bioassay-guided isolation from natural sources. In reverse chemical ecology, active compounds are identified first by their interactions with olfactory proteins, such as odorant receptors (ORs), and subsequent behavioral assays [2].

Semiochemicals must first activate receptors housed in olfactory neurons in the peripheral sensory system of insects (e.g., antennae, maxillary palps, and proboscis) [3] to elicit behavior. Highly expressed odorant receptors are intuitive targets for reverse chemical ecology because high expression levels imply that they are involved in the reception of crucial semiochemicals. Therefore, de-orphanizing highly expressed ORs may lead to the discovery of novel attractants or repellents, provided the test ORs are adequately expressed and challenged with appropriate compounds.

The two-electrode voltage-clamp (TEVC) recording system has been widely used to de-orphanize ORs [4, 5]. With this technique, a test OR is co-expressed with the co-receptor Orco in *Xenopus laevis* oocytes and challenged with test compounds. Co-expression is necessary because, in more evolved insect species, the binding units of ORs form ion channels (pathways) with the obligatory odorant receptor co-receptor Orco. In more primitive insects, tetramers of the binding unit *per se* form ion channels [6]. Previously, we have de-orphanized ORs in the genome of the Southern house mosquito, *Culex quinquefasciatus* [7–12], as part of our reverse chemical ecology approach. For example, we expressed CquiOR36/CquiOrco in *Xenopus* oocytes, challenged them with a panel of 230 odorants, and identified acetaldehyde as the most potent ligand [13]. Subsequent behavioral tests showed that acetaldehyde is an oviposition attractant.

Considering the metabolic resources used for their synthesis, it is conceivable that highly expressed ORs play crucial roles in the ecology of the insect. Intriguingly, however, many ORs from the Southern house mosquito were found to be “silent,” i.e., no compounds in our extensive panel of odorants elicit measurable currents in these CquiORx/CquiOrco-expressing oocytes. One such case is CquiOR125, which is the second-most expressed OR in *Cx. quinquefasciatus* antennae [9, 13, 14]. We surmised that testing an ortholog from the yellow fever mosquito, *Aedes aegypti*, AaegOR11 [15], might shed some light on the function of the highly expressed CquiOR125. AaegOR84, AaegOR113, and AaegOR11 are three of the most expressed ORs in the female yellow fever mosquito antennae [15]. While AaegOR84 and AaegOR113 have no orthologs in the *Cx. quinquefasciatus* genome, AaegOR11 shares 73% amino acid identity with CquiOR125.

We cloned these three ORs and challenged AegORx/AaegOrco-expressing oocytes with a panel of 286 compounds. AaegOR84/AaegOrco- and AaegOR113/AaegOrco-expressing oocytes were not activated by any compound in our panel of odorants but elicited small inward currents when challenged with the Orco ligand candidate, OLC 12 [16]. By contrast, AaegOR11/AaegOrco-expressing oocytes generated strong inward currents when stimulated with fenchone, 2,3-dimethylphenol, and other compounds. Interestingly, AaegOR11 does not discriminate fenchone stereoisomers. In-door and field tests suggest that fenchone is neither an oviposition attractant nor a repellent for the yellow fever mosquito, and 2,3-dimethylphenol is a potent repellent.

## Results and Discussion

### Cloning and de-orphanization of ORs

We cloned AaegOR11, 84, and 113 using cDNA obtained from *Ae. aegypti* mosquitoes collected in Clovis, CA, as a template. Four clones of AaegOR11 gave identical sequences (GenBank OM568708), but they differed in five residues compared to the AaegOR11 amino acid sequence in VectorBase (AAEL011583). Specifically, Ser163Thr, Met167Ile, Asn240His, Met249Ile, and Met338Ile. In short, three methionine residues in the VectoBase sequence are Ile in the Clovis strain, a His replaces an Asn, and a Ser changed to a Thr. Our AaegOR84 and AaegO113 clones showed sequences identical to those in VectorBase, i.e., AAEL017043 and AAEL017123, respectively.

We co-expressed each OR with the odorant receptor co-receptor Orco in *Xenopus* oocytes and challenged them with a large panel of odorants. AaegOR113/AaegOrco-expressing oocytes did not respond to any test compound, except for a small current elicited by the Orco ligand OLC 12 [16]. Likewise, no currents were recorded when AaegOR84/AaegOrco-expressing oocytes were challenged with individual compounds in our panel of odorants, except for OLC 12-induced currents. By contrast, AaegOR11/AaegOrco-expressing oocytes responded to twelve compounds in our panel, specifically, fenchone, 2,3-dimethylphenol, 2,6-dimethylphenol, 3,4-dimethylphenol, 2-methylphenol, 4-methylcyclohexanol, acetophenone, guiacol, 2-acetylthiophene, indole, 1,2-dimethoxybenzene, and (*Z*)-3-hexenyl acetate. We then challenged AaegOR11/AaegOrco-expressing oocytes at least six times with each active compound (Fig 1). Fenchone and 2,3-dimetylphenol, 3,4-dimethylphenol, 4-methylcyclohexanol, and acetophenone gave the most robust responses (Fig 1).

**Fig 1.**
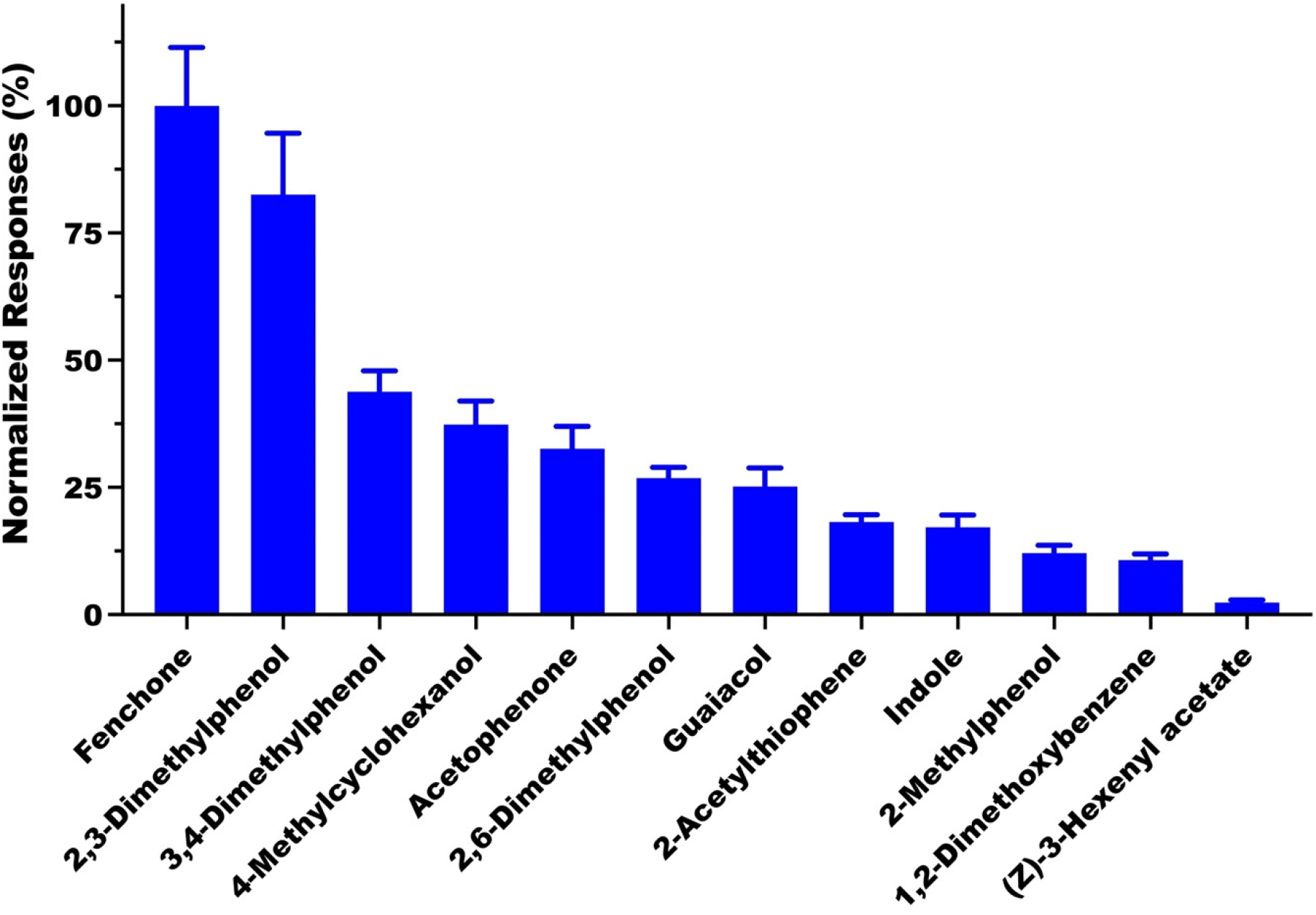
Quantification of current responses recorded from oocytes co-expressing AaegOR11 and AaegOrco. All compounds were tested at 1 mM (source dose, n = 6). For clarity, bars are displayed in decreasing order of response (mean ± SEM).

Concentration-dependent relationships for the five ligands that produced the most robust responses confirm that fenchone was the most potent ligand, followed by 2,3-dimethylphenol, and 3,4-dimethylphenol (Fig 2).

**Fig 2.**
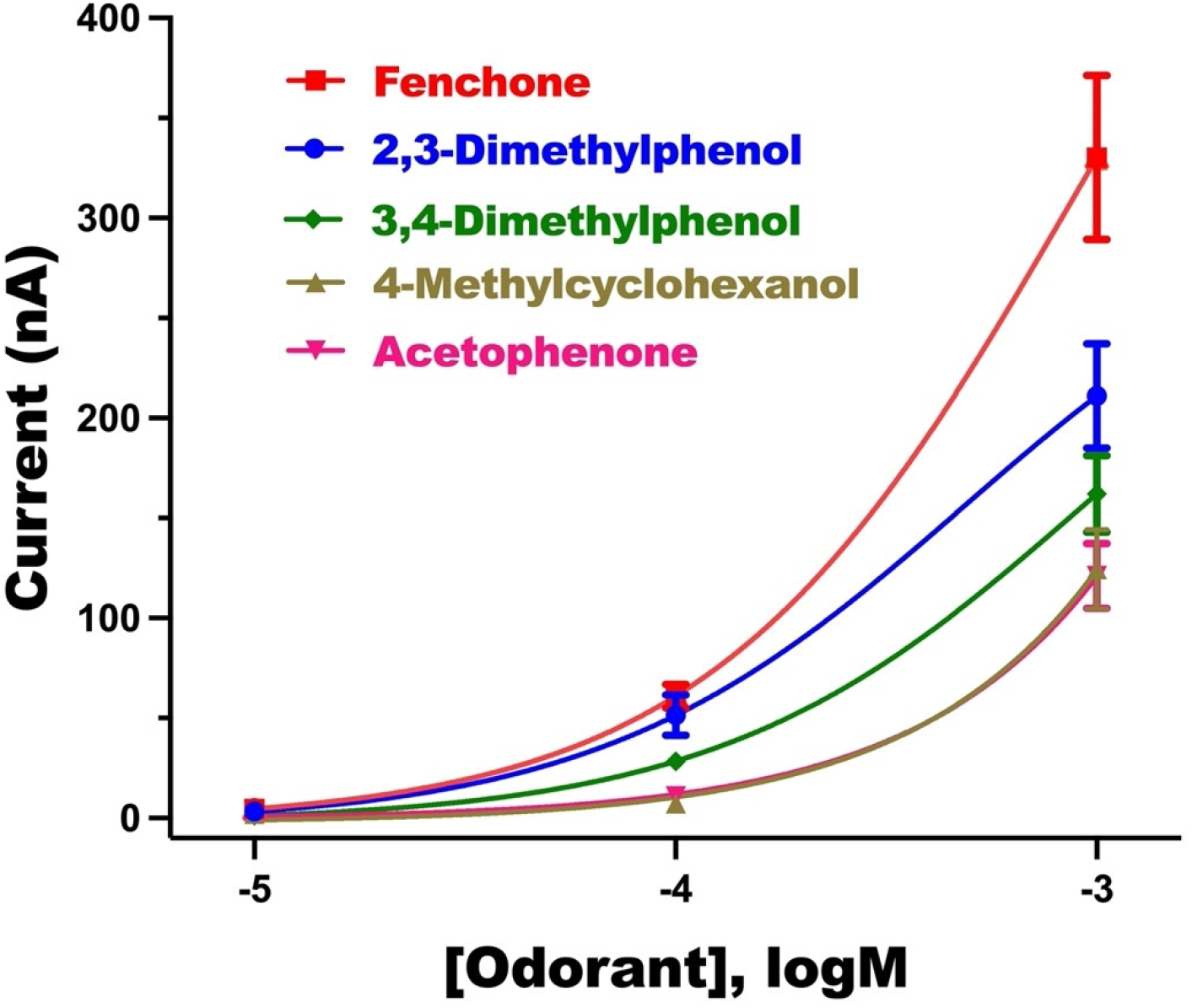
Concentration-dependent responses were recorded from AaegOR11/AaegOrco-expressing oocytes. Currents were recorded from seven oocytes challenged at 0.01, 0.1, and 1 mM, with seven replicates (n = 7) per stimulus per dose.

We then asked whether this receptor could discriminate fenchone stereoisomers. Interestingly, AaegOR11/AaegOrco-expressing oocytes responded equally to (1R,4S)-fenchone (=(-)- fenchone, PubChem CID #82229) and (1S,4R)-fenchone (=(+)-fenchone, PubChem CID #1201521) (Fig 3A). A dose-dependent curve confirmed that the receptor responds to the two stereoisomers equally (Fig 3B)

**Fig 3.**
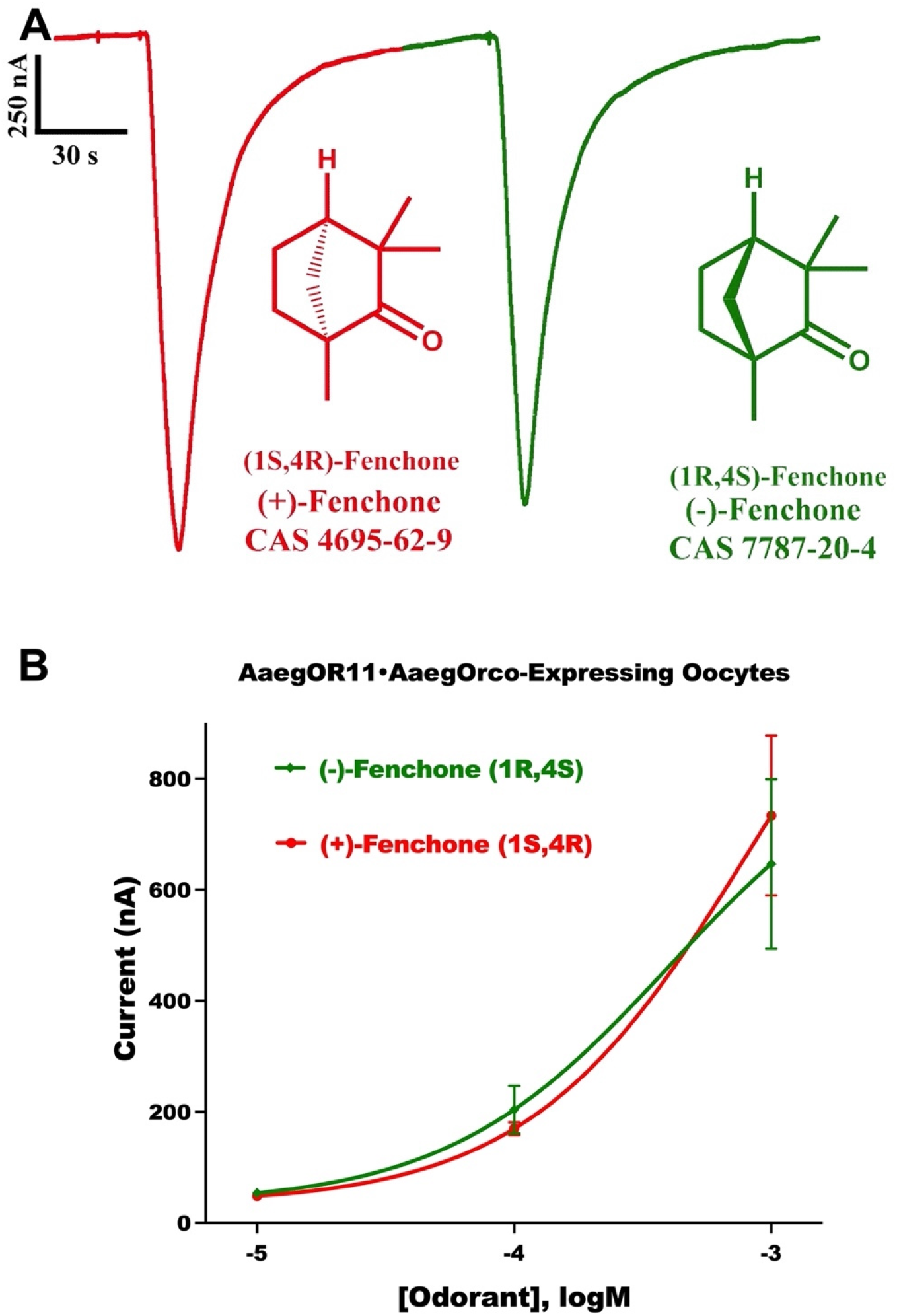
Responses were recorded from oocytes co-expressing AaegOR11 and AaegOrco. (A) A representative trace with recordings from the same oocyte challenged with (+)- and (-)- fenchone at 1 mM. Similar results were obtained when oocytes were stimulated first with (-)- and then (+)-fenchone. (B) Concentration-dependent relationships obtained with oocytes (n = 3) challenged with the two isomers at three different doses (n = 6-9 replicates/dose).

### Functional studies

(+)-Fenchone has been reported as a potent repellent against *Ae. aegypti* [17] and *Ae. albopictus* [18] comparable to DEET, but a moderate repellent 30 min after sample preparation and about 50% activity one hour later [17]. We tested both (+)-fenchone and (-)-fenchone with our surface-landing and feeding assay [9, 19, 20], and using freshly prepared samples. In our hands, neither (+)-fenchone, nor (-)-fenchone showed significant repellency activity against *Ae. aeypti* (Fig 4).

**Fig 4.**
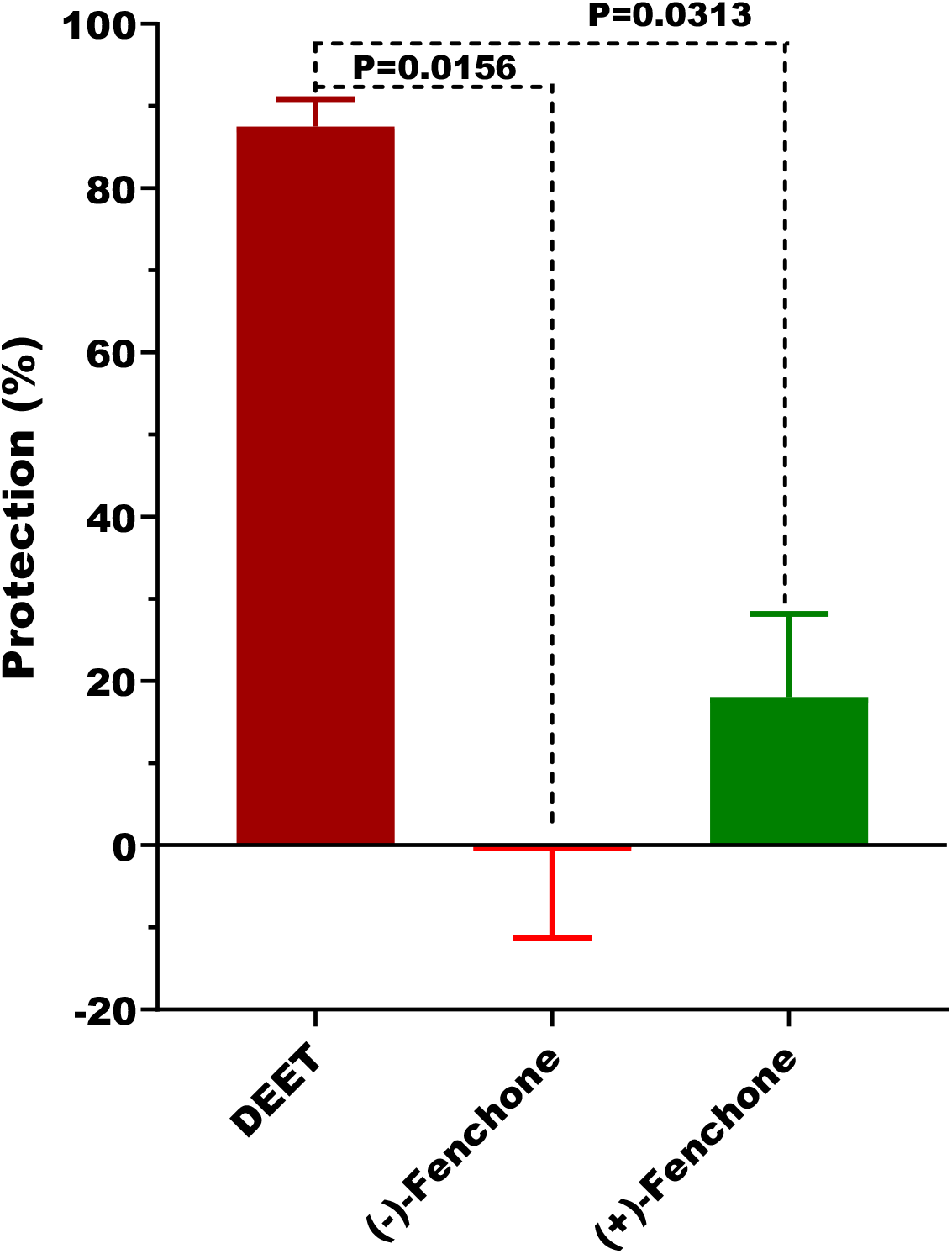
Fenchone repellency measured in the surface-landing and feeding assay. Repellency is expressed in percent protection for DEET (n = 7), (-)-fenchone (n = 8) and (+)- fenchone (n = 6). Tests were conducted in tandem by measuring the responses to DEET (2 replicates, one on each side of the arena) and two replicates of (+)- or (-)-fenchone and repeating this cycle until the response to fresh samples of DEET started to decrease. Data were analyzed by the Wilcoxon matched-pairs signed rank test

The (+)-fenchone sample reported being a potent repellent against *Ae. aegypti* was extracted and isolated from fruits of the common fennel *Foeniculum vulgare* Miller [17]. The reported specific rotation [17] suggests an enantiopure compound was used in the previous work. In any case, we analyzed our sample to rule out a possible problem with enantiomeric impurities. Gas chromatography analysis by separation on a capillary column with a chiral stationary phase showed that our (+)-fenchone sample was enantiopure, whereas the (-)-fenchone sample contained a small percent of (+)-fenchone (Fig 5).

**Fig 5.**
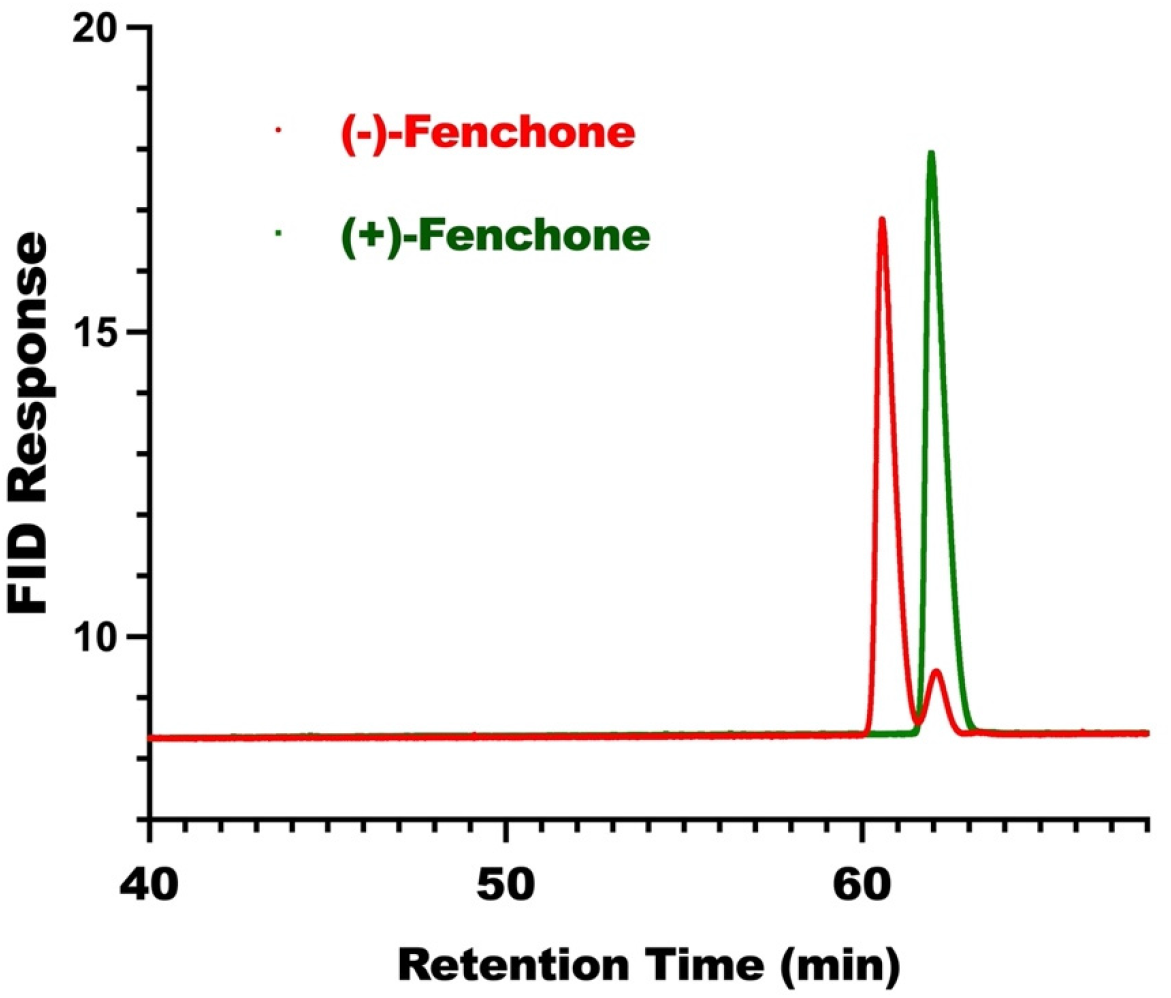
Separation of fenchone isomers on a gas chromatograph equipped with a capillary column with a chiral stationary phase. Baseline separation of (-)- and (+)-fenchone was achieved with an HP-CHIRAL-20B column. For clarity, the traces were reconstructed by Prism using the FID data imported from the Agilent software.

Given the discrepancy between the literature [17, 18] and our findings, we tested whether the (+)- and (-)-fenchone repel *Cx. quinquefasciatus*, but neither isomer showed repellency activity. The protection elicited by (+)-fenchone at 1% was −52±27.5% (n = 8), whereas DEET showed 92.2±2.5% (n = 4). Similarly, (-)-fenchone gave low protection −18±20.2% (n = 8); DEET, 100% (n = 4). In unusual cases [21], semiochemicals are active only when presented as a racemic mixture. We, therefore, tested a (+/-) mixture of fenchone and observed no activity: 9.5±8.0 % protection (n = 8); DEET 90.1±2.4% (n = 8).

Next, we tested whether fenchone could act as an oviposition attractant for the yellow fever mosquito. First, we conducted in-door assays and focused on (-)-fenchone at doses ranging from 10^−2^ % to 10^−7^% (Fig 6A). For each dose, 100 μl of a concentrated fenchone solution in ethanol was added to a cup filled with 100 ml of tap water. One hundred μl of ethanol was added to each control cup with 100 ml tap water. There were no significant differences in the number of eggs laid in treatments at doses lower than 0.01% (10^−2^ %) compared with their respective controls. However, the number of eggs laid in the 0.01% (-)-fenchone cups was significantly higher than those laid in control cups (P=0.0244, n = 6, two-tailed, paired t-test). Considering that activity was observed at the highest dose tested, we repeated these experiments with 4 doses, 0.001, 0.005, 0.01, and 0.05% (Fig 6B). We found no significant difference between treatment and control in all doses tested, including 0.01% (10^−2^ %).

**Fig 6.**
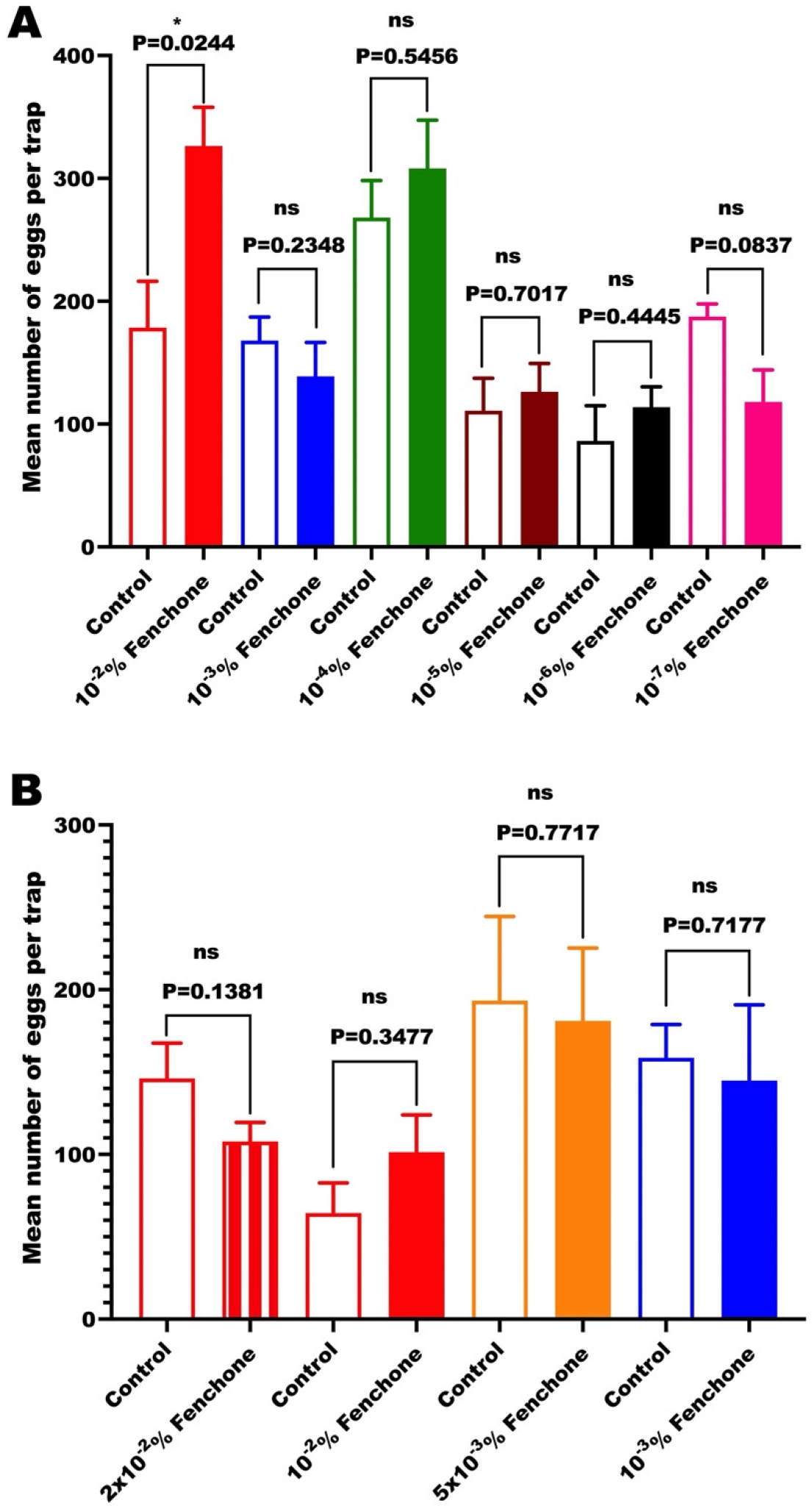
Results of in-door oviposition assays with fenchone at various doses. (A) The first screening of (-)-fenchone activity at multiple doses, and (B) a closer examination of the responses to concentrations closer to 10^−2^%. All tests were performed with six replicates (n = 6 per dose). Data passed the Shapiro-Wilk normality test and were, therefore, analyzed by paired, two-tailed t-test.

Given this discrepancy in the two in-door trials regarding the activity of (-)-fenchone at 10^−2^ % (Fig 6), we conducted a third trial with a single dose of 0.01% that included a larger number of replicates (Fig 7). Again, there was no significant difference between treatment and control (P=0.1882, n = 22, two-tailed paired t-test).

**Fig 7.**
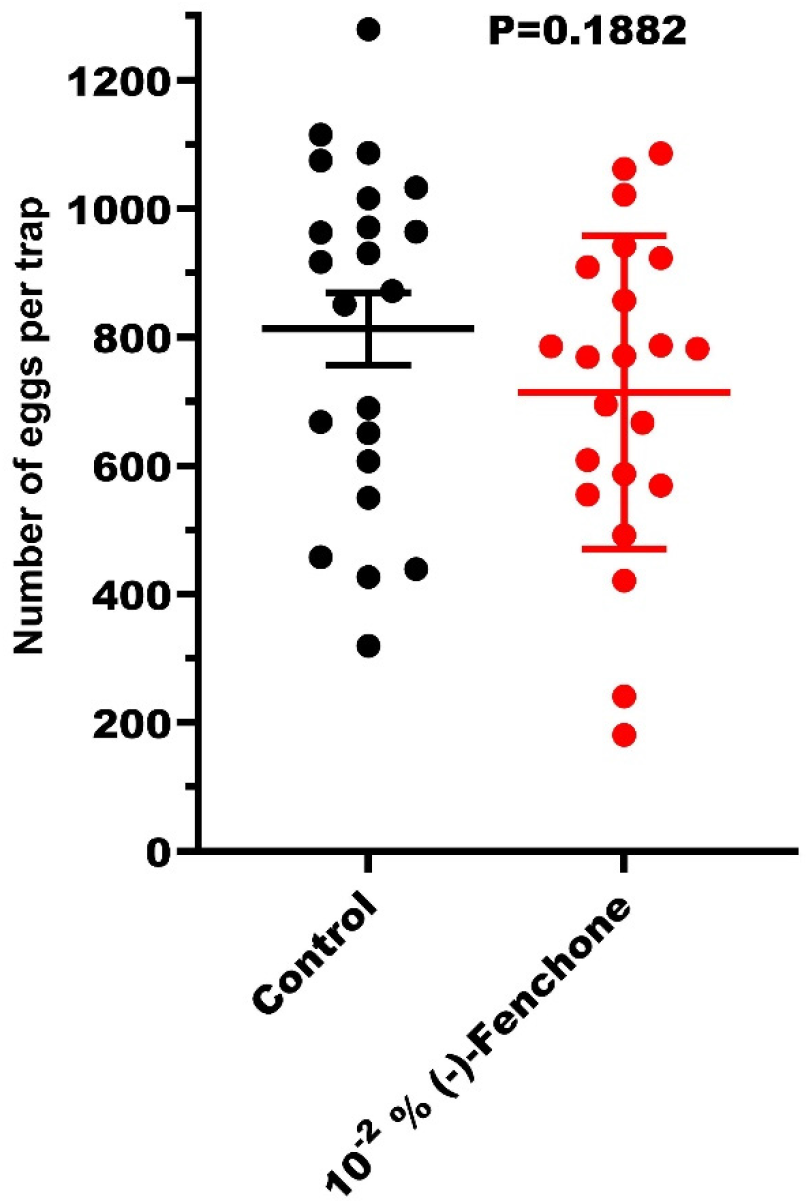
Evaluation of fenchone as a putative oviposition attractant in a single dose. Each of the six cages used had control and treatment; the experiment was replicated four times. Two outliers were omitted for clarity (cage 2: control, 731; treatment, 1477; cage 3: control, 1952; treatment, 959). The complete dataset, including the two outliers, did not pass the Shapiro-Wilk normality test and was analyzed by the Wilcoxon matched-pairs signed rank test (p=0.2768, n = 24).

Lastly, we performed field tests comparing the deposition of eggs in cups containing Orchard grass infusion [22] (positive control), racemic fenchone, or water (control) (Fig 8). Traps were placed in seven properties in Reedley (Fresno County, California), with three replicates of one week each. The total number of eggs laid in control, infusion, and fenchone cups, respectively, were: 224, 741, and 259 (week 1); 297, 797, and 546 (week 2), and 276, 984, and 567 (week 3). While cups loaded with infusion had significantly more eggs than control cups (145.7±22.7 vs. 44.3±7.9; adjusted P=0.0002, n = 18, one-way ANOVA), egg counts in cups holding racemic fenchone did not significantly differ from egg counts in control traps (77.3±16.7 vs. 44.3±7.9; adjusted P=0.2969, n = 18, one-way ANOVA). This led us to conclude that fenchone solution was no more or less attractive than water but confirmed that grass infusion offered higher egg-laying attractive properties. For now, mosquito control agencies should still have cups filled with Orchard grass infusion for *Ae. aegypti* oviposition field surveillance purposes.

**Fig 8.**
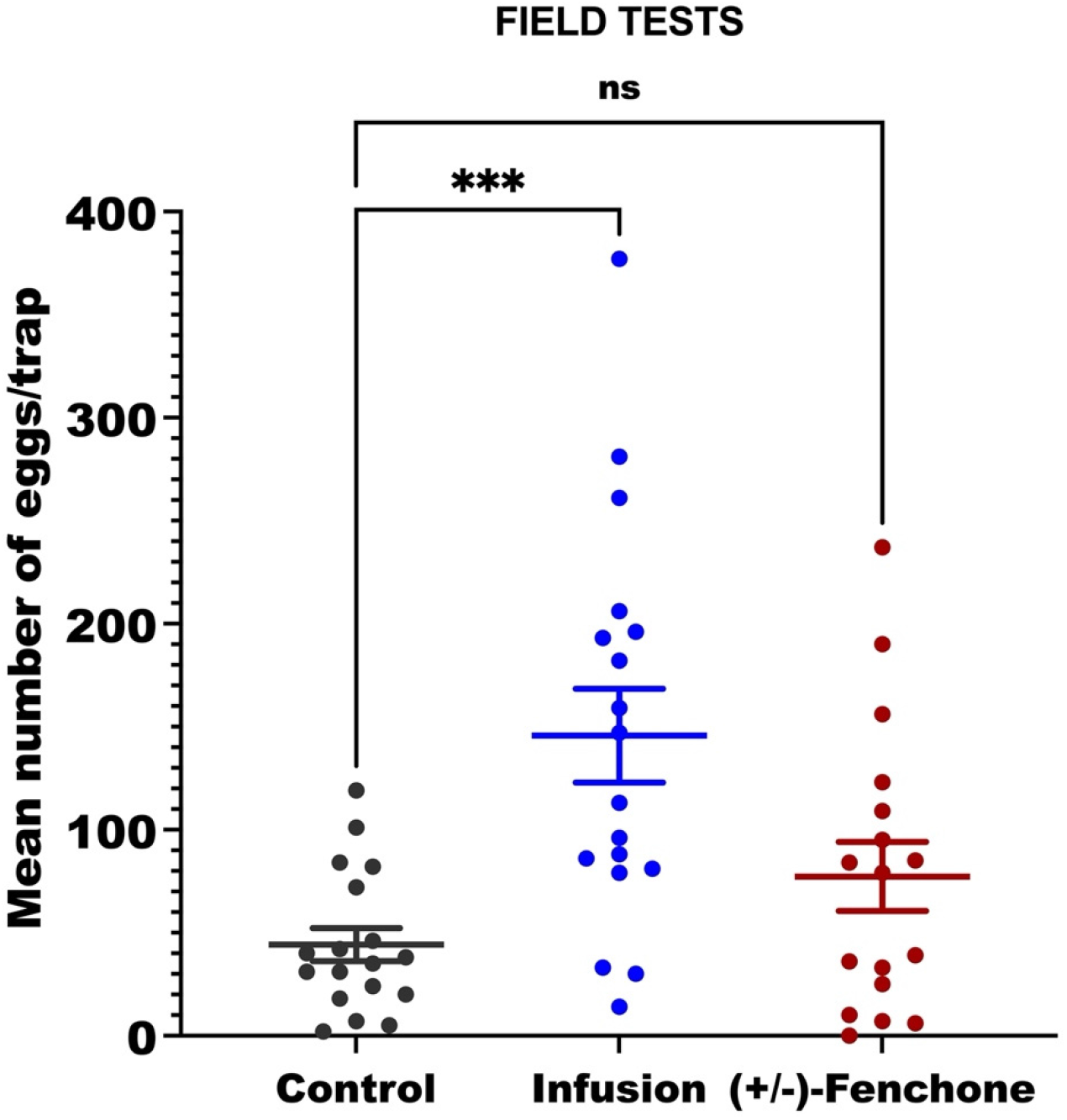
Results of field tests in Reedley (Fresno Co.) in late summer. Control traps were baited with a 7-day-old Orchard grass (*Dactylis glomerata* L.)-based infusion; fenchone traps were filled with 0.01% racemic fenchone. A set of control-, infusion- and fenchone-trap, separated by 30 m, was placed in seven properties in Reedley. Eggs were counted after seven days. These experiments were repeated three times late in summer with average maximal and minimal temperatures of 37 and 19°C, respectively.

Surprisingly, the most potent ligand for a highly expressed receptor did not show any activity as either a repellent or an oviposition attractant. We cannot rule out that fenchone plays a crucial role in the chemical ecology of the yellow fever mosquito, particularly for behaviors not measured in our study, including, but not limited to, mating behavior, attraction to flowers, and pollination. Additionally, fenchone may elicit repellency or oviposition attraction in synergy with other semiochemicals. However, our studies demonstrate that fenchone is neither a repellent nor an oviposition attractant when used as a single compound.

Lastly, we tested whether the second most potent ligand, 2,3-dimethylphenol, would repel mosquitoes. In contrast to fenchone, 2,3-dimethylphenol at 1% showed significantly stronger (P = 0.0133, Wilcoxon matched-pairs signed rank test) repellency activity against *Cx. quinquefasciatus* mosquitoes: 94.3±0.8 % protection (n = 32), than DEET at 1%: 81.5±2.6 % (n = 22). These findings suggest that AaegOR11, one of the ORs most expressed in the female yellow fever mosquito antennae [15], is involved in the reception of repellents.

AaegOR11 is 73% identical to CquiOR125. Of note, there is a proposition under consideration to renaming CquiOR125 to CquiOR11 (Dr. Carolyn McBride, personal communication). Previous work showed that none of the ligands for AaegOR11 identified in this study activated CquiOR125 in a previous study [9]. Indeed, CquiOR125 did not respond to any compound in a large panel of odorants [9]. We compared the predicted structures of these two receptors and did not find any structural features that might explain CquiOR125 lack of activation.

Recently, the predicted structures of 200 million proteins have been reported [23, 24]. Additionally, two experimental structures have been reported to date, the cryo-EM structure of the co-receptor Orco from the parasitic fig wasp *Apocrypta bakeri*, AbakOrco, and the cryo-EM OR structure from the jumping bristletail, *Machilis hrabei*, MhraOR5 [6].

The AlphaFold model for AaegOR11, predicted by DeepMind (A0A1S4FTS1) (Fig 9A) and MhraOR5 have structurally homologous cores (Fig 9B,C). The room mean-square deviation, RMSD, between 155 pruned atom pairs was 1.18 Å (and 4.5 Å across all 368 pairs), as calculated by Chimera [25]. The transmembrane (TM) domains, which form the ligand binding pocket [6], are shifted in AaegOR11 relative to MhraOR5-eugenol (Fig 9B,C). AaegOR11 TM2 closer to the eugenol binding pocket in MhraOR5 (inward shift), whereas M3, M4, and M5 moved away from the eugenol binding pocket in MhraOR5 (outward shift). The C-terminus of TM-3 with a peptide (S-166-MDSF-170) protrudes beyond the predicted extracellular domain (Fig. 9A,C). Of note, the amino acid sequence of the AaegOR11, which we cloned from *Ae. aegypti* mosquitoes from Clovis, differs from the sequence in VectorBase in the following residues: Ser163Thr (TM-3), Met167Ile (TM-3), Asn240His (intracellular segment-2), Met249Ile (intracellular loop-2), and Met338Ile (TM-6) (Fig 9D). These mutations are far from the predicted binding pocket (Fig 9D).

**Fig 9.**
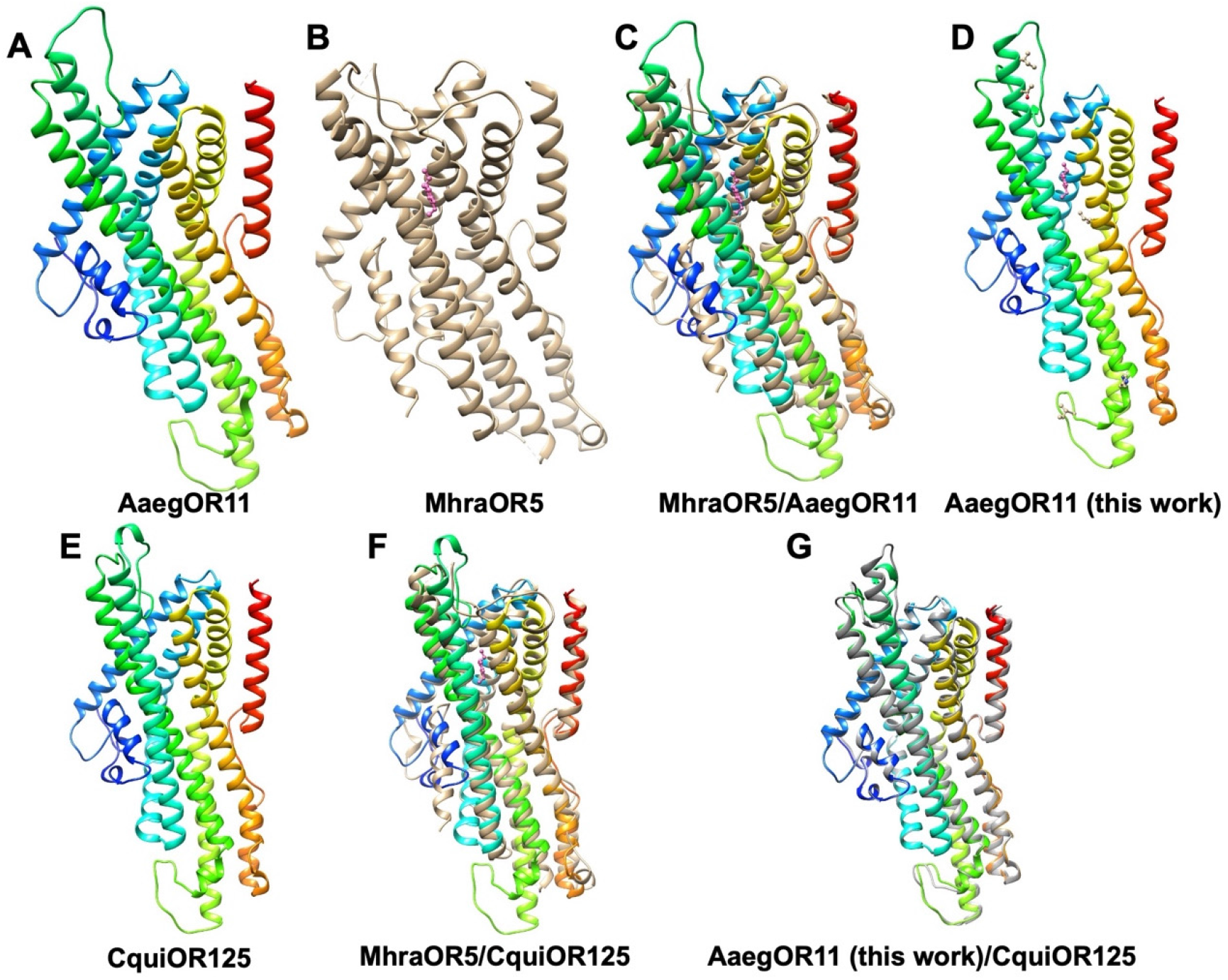
AlphaFold models for mosquito ORs compared to the cryo-EM MhraOR5/eugenol structure. (A) AegOR11 model (A0A1S4FTS1) based on the primary sequence in VectorBase (AAEL011583). (B) MhraOR5-eugenol experimental structure [26]; eugenol is displayed in hot pink ball and stick. (C) Superposed structures in (A) and (B). (D) Predicted model for AaegOR11 based on the primary sequence reported in this study (OM568708). (E) AlphaFold model (B0X7M5) for CquiOR125 (CPIJ015178). (F) Superposed CquiOR125 model with the experimental MhraOR5-eugenol structure (in tan; eugenol in hot pink). (G) Superposed models of AaegOR11 (predicted based on the amino acid sequence reported here, in dark grey) and CquiOR125. Unless otherwise specified, subunits and loops were colored in a rainbow pattern from the N-terminus (purple) to the C-terminus (red).

The AlphaFold model structure (B0X7M5) for CquiOR125 (CPIJ015178) suggests a homologous structure to MhraOR5 (Fig 9E, 9F). The RMSD between 169 pruned atom pairs was 1. 1 Å, and across all 364 pairs, 4.2 Å. No apparent structural feature in this model would suggest a non-functional OR. By contrast, comparing another non-functional OR, CquiOR55 (CPIJ005401) [9], with MhraOR5 provides a reasonable explanation for the lack of activity. For example, the ion pathway-forming C-terminus [6] is missing in CquiOR55 (B0WDQ3). CquiOR125 and AaegOR11 are structurally homologous. The RMSD for AaegOR11 (model based on OM568708 primary sequence) and CquiOR125 (model based on CPIJ005401 amino acid sequence) were 0.8 Å between 378 pruned atom pairs and 1.6 Å across all 414 pairs.

In summary, we have de-orphanized an OR highly expressed in the antennae of the female yellow fever mosquito, AaegOR11, which shared high amino acid identity with CquiOR125, a “silent” receptor from the southern house mosquito. AlphaFold models showed that both CquiOR125 and AaegOR11 have structural cores homolog to those in the cryo-EM structure of MhraOR5. In the *Xenopus* oocyte recording system, AaegOR11, co-expressed with AaegOrco, generated robust responses when challenged with fenchone, 2,3-dimethylphenol, 3,4-dimethylphenol, 4-methylcyclohexanol, and acetophenone. Fenchone did not elicit blood-feeding repellency or oviposition activity. By contrast, behavioral measurement showed that 2,3-dimethylphenol is a potent blood-feeding repellent. Considering that AaegOR11 is a repellent-detecting OR, it is conceivable that AaegOR11 orthologue in the genome of *Cx. quinquefasciatus*, CquiOR125, is also a “repellent receptor,” but the active ligand is still elusive.

## Methods

### Mosquitoes

*Culex quinquefasciatus* mosquitoes used in this study originated from the CQ1 colony, established in the 1950s, from specimens collected in Merced County, CA. The CQ1 colony has been used as a standard insecticide susceptible strain in insecticide bioassays [27]. In Davis, mosquitoes were maintained at 27±1°C, 75±5% relative humidity, and under a photoperiod of 12:12 h.

*Aedes aegypti* mosquitoes used for molecular studies and blood feeding repellency choice assays originated from second filial generation individuals originating from adults collected in Clovis, (Fresno County, CA). The first two of the three laboratory oviposition repellency/attractiveness trials were conducted using females reared from *Ae. aegypti* eggs that were laid by wild females in oviposition cups placed at multiple properties in Reedley. In the third repellency/attractiveness trial, females reared from the eggs laid by females in the first two trials were used.

### Cloning and de-orphanization

The p-GEMHE plasmids for AaegOR11, AaegOR84, and AaegOR113 were obtained by using *Ae. aegypti* female antennae cDNA [28] as a template and the infusion primers below. The resulting PCR products were purified with gel electrophoresis, QIAquick gel extraction kit (Qiagen, Germantown, MD), and then cloned into pGEMHE using In-Fusion HD Cloning Kit according to the manufacturer’s instructions (Clontech).

PCR In-Fusion primers were designed according to the user manual.
InFuAaOR11F primer:
5’- GATCAATTCCCCGGGACCATGCAGCTGAAAGACGAATGG −3’ and
InFuAaOR11R primer:
5’- CAAGCTTGCTCTAGATTAGCCGGCAGCTTGTTTC −3’;
InFuAaOR84F primer:
5’- GATCAATTCCCCGGGCACATCATGAACGCGATTGT −3’ and
InFuAaOR84R primer:
5’- CAAGCTTGCTCTAGAGCCATTGCATTATTCCGATT −3’.
InFuAaOR113F primer:
5’- GATCAATTCCCCGGGACCATGTTTGCGGAAATTCGTGGC −3’ and
InFuAaOR113R primer:
5’- CAAGCTTGCTCTAGACTAAACCATATTGATCAGAAATGTTAACACTG −3’.

The colonies from transformations were verified by regular PCR. The positive clones were cultured and subsequently extracted using the QIAprep Spin Miniprep kit (Qiagen) and sequenced by ABI 3730 automated DNA sequencer.

TEVC was performed as previously described [19]. In brief, linearized pGEMHE-AaegORs were used as templates to transcribe into capped cRNA with poly(A) using an mMESSAGE mMACHINE T7 kit (Ambion, Austin, TX) following the manufacturer’s protocol. The cRNAs were dissolved in RNase-free water and adjusted to a concentration of 200 μg/mL by UV spectrophotometry (NanoDrop™ Lite Spectrophotometer). 9.2 nl of a 50/50 mixture of a test OR and AaegOrco [25] cRNAs were microinjected into V or VI *Xenopus* oocytes (purchased from EcoCyte Bioscience, Austin, TX). Then, injected oocytes were incubated at 18°C for 3–7 days in modified Barth’s solution [in mM: 88 NaCl, 1 KCl, 2.4 NaHCO_3_, 0.82 MgSO_4_, 0.33 Ca(NO_3_)_2_, 0.41 CaCl2, 10 HEPES, pH 7.4] supplemented with 10 μg/mL of gentamycin, 10 μg/mL of streptomycin, and 1.8 mM sodium pyruvate. Test oocytes were placed in a perfusion chamber and challenged with a panel of odorants (see below). Currents were amplified with an OC-725C amplifier (Warner Instruments, Hamden, CT) holding the voltage at −80 mV and a low-pass filter at 50 Hz, and digitized at 1 kHz. Data acquisition and analysis were carried out with Digidata 1440A and pClamp10 software (Molecular Devices, LLC, Sunnyvale, CA). To standardize the measurement, 6-7 days old eggs were used for TEVC measurement.

### Panel of odorants

Oocytes were challenged with the following compounds: 1-butanol, 1-pentanol, 1-hexanol, 1-heptanol, 1-octanol, 1-nonanol, 2,3-butanediol, 2-butoxyethanol, 3-methyl-1-butanol, 2-hexen-1-ol, 3-hexen-1-ol, 1-hexen-3-ol, 1-hepten-3-ol, 3-octanol, 1-octen-3-ol, 2-octanol, 2-butanol, 2-nonen-1-ol, 2-pentanol, 4-methylcyclohexanol, 1-hexadecanol, menthyl acetate, methyl acetate, ethyl acetate, propyl acetate, butyl acetate, pentylacetate, hexyl acetate, heptyl acetate, octyl acetate, nonyl acetate, decyl acetate, methyl propionate, ethyl propionate, methyl butyrate, ethyl butanoate, methyl hexanoate, ethyl hexanoate, ethyl 3-hydroxyhexanoate, ethyl 3-hydroxybutanoate, (*E*)-2-hexenyl acetate, (*Z*)-3-hexenyl acetate, α-terpinene, γ-terpinene, ethyl lactate, methylsalicylate, 1-octen-3-yl acetate, isopentyl acetate, m-tolyl acetate, ethyl phenylacetate, geranyl acetate, octadecyl acetate, propanal, butanal, penatanal, hexanal, (*E*)-2-methyl-2-butenal, heptanal, octanal, nonanal, decanal, undecanal, 1-dodecanal, (*E*)-2-hexenal, (Z)-8-undecenal, (*E*)-2-heptenal, (*E*)-2-nonenal, phenylacetaldehyde, furfural, 2-butanone, 2-heptanone, geranylacetone, 6-methyl-5-hepten-2-one, 5-methyl-2-hexanone, 2,3-butanedione, 3-hydroxy-2-butanone, 2-undecanone, (-)-menthone, 2-tridecanone, 2-nonanone, 5-isobutyl-2,3-dimethylpyrazine, (*S*)-(-)-perillaldehyde, (1 *S*,4*R*)-fenchone, (1*R*,4*S*)-fenchone, cyclohexanone, acetophenone, (α+β)-thujone, p-coumaric acid, isovaleric acid, 1-dodecanol, dodecanoic acid, (±)-lactic acid, ethanoic acid, propanoic acid, butanoic acid, isobutyric acid, 2-oxobutyric acid, pentanoic acid, 2-oxovaleric acid, palmitoleic acid, oleic acid, hexanoic acid, (*E*)-2-hexenoic acid, 5-hexanoic acid, (*E*)-3-hexenoic acid, heptanoic acid, octanoic acid, nonanoic acid, decanoic acid, trimethylamine, n-tridecanoic acid, linoleic acid, ammonia, trimethylamine, propylamine, butylamine, pentylamine, hexylamine, heptylamine, camphor, octylamine, 1,4-diaminobutane, cadaverine, 1,5-diaminopentane, benzaldehyde, phenol, 2-methylphenol, 3-methylphenol, 4-methylphenol, 4-ethylphenol, 3,5-dimethylphenol, 2,3-dimethylphenol, guaiacol, 2-methoxy-4-propylphenol, 2-phenoxyethanol, 1,2-dimethoxybenzene, benzyl alcohol, 2-phenylethanol, 1-phenylethanol, phenyl ether, isoprene, limonene, linalyl acetate, α-humulene, linalool oxide, geraniol, nerol, thymol, (±)- linalool, eucalyptol, citral, eugenol, α-pinene, ocimene, (±)-citronellal, indole, permethrin, acetaldehyde, 3-methylindole, 3-pentanol, 3-methyl-2-butanol, 3-methyl-2-buten-1-ol, 2-methyl-3-buten-2-ol, γ-valerolactone, γ-hexalactone, γ-octalactone, γ-decalactone, α-phellandrene, nerolidol, γ-dodecalactone, jasmine, 1-octen-3-one, dibutyl phthalate, dimethyl phthalate, menthol, benzyl formate, 2-acetylthiophene, phenethyl formate, carvone, isovaleraldehyde, n-methylbenzamide, 3-methylbenzamide, ethyl stearate, tetradecanoic acid, methyl myristate, p-cymene, sabinene, 2-tridecanone, terpinolene, valencene, 2,4-pentanedione, (*R*)-(+)-pulegone, β-myrcene, 5-isobutyl-2,3-dimethylpyrazine, phenyl propanoate, phenyl isobutyrate, eugenyl acetate, (+)-δ-cadinene, (+)-limonene oxide, 2-ethyltoluol, 2,4-hexadienal, 2-methyl-2-thiazoline, phenethyl propionate, ethyl (2*E*,4*Z*)-decadienoate, methyl anthranilate, α-hexylcinnamaldehyde, 2-pentanone, 2-hexanone, 2-octanone, 4,5-dimethylthiazole, (*E,E*)-farnesol, (*E,E*)-farnesyl acetate, farnesene, 4-dimethylamino-1-naphthaldehyde, α-methylcinnamaldehyde, cinnamyl alcohol, α-terpineol, citronellol, (*E*)-cinnamaldehyde, (-)-caryophyllene oxide, β-caryophyllene, carvacrol, terpinen-4-ol, neo-alloocimene, palmitic acid methyl ester, 2-ethylhexanoic acid, pentadecanoic acid, geranyl isovalerate, geranyl butyrate, α-cetone, 7-hydroxcitronellal, isopropyl myristate, methyl N,N-dimethylanthranilate, pyridine, pyrrolidine, pyrrolidinone, amiloride, allethrin, allopurinol, methyl-p-benzoquinone, lilial, styrene, indole-3-aldehyde, dimethyl trisulfide, methyl disulfide, cyclohexanol, 2-methylbutyl 2-methylbutanoate, sabinene, citronellyl acetate, methyl geranate, 3-methylbutyl 2-methylbutanoate, *tert*-butanol, 2,4-dimethylphenol, 2,5-dimethylphenol, 2,6-dimethylphenol, 3,4-dimethylphenol, ethyl undecanoate, neryl acetate, isobutyl acetate, methyl sulfide, 2-hexanol, (+)-3-carene, ethyl butyrate, α-ionone, β-ionone, formic acid, linalyl formate, diethyl sebacate, 3,5-di-*tert*-butyl-4-hydroxyacetophenone, 2,4-di-*tert*-butylphenol, 2-nonanol, N-methylpiperidine, 5-ehtyl-2-methylpyridine, piperine, 1-methylpiperidine, 2-methylpiperidine, 3-methylpiperidine, 4-methylpiperidine, dimethyl carbonate, 5-ethyl-2-methylpyridine, 3-pentyn-1-ol, 4-methylthiazole, 4,5-dimethylthiazole, tryptamine, dicyclohexyl disulfide, and isoamyl alcohol. The Orco ligand candidate 2-{[4-Ethyl-5-(4-pyridinyl)-4*H*-1,2,4-triazol-3-yl]sulfanyl}-*N*-(4-isopropylphenyl)acetamide (OLC 12 [16] = VUAA-3 [29]) was used for confirming receptor protein expression.

### Chemical analysis

Gas chromatography analyses were performed on an Agilent 6890 Series GC System equipped with a flame ionization detector (FID) and a chiral capillary column (HP-Chiral-20B, 30 m x 0.25 mm; 0.25 μm). The injection port and the FID detector were operated at 250°C. The oven temperature was set at 85°C for 70 min, with a 5 min post-run at 210°C.

### Repellent assays

Repellency was measured using the surface landing and feeding assays, previously described in detail [9, 20]. In short, two Duddle bubbling tubes were inserted inside a mosquito cage (arena) through a wooden board attached to the back of the cage. The tubes were covered with non-lubricated condoms to avoid contamination; they were replaced every five tests [19]. Water circulated inside the tubes to maintain the surface at 37°C. A needle was inserted through the wooden board above each tube to deliver CO_2_ (50 ml/min). A dental cotton roll holding defibrinated sheep blood (100 μl) was inserted between a Duddle tube and a needle. One filter paper cylinder was impregnated with solvent only (control; 200 μl of hexane) and the other with a solution of the test compound (200 μl of a 1% solution in hexane). After the solvent evaporated, the cylinders were placed around each tube and held in place by insect pins. Repellency was measured using cages with one hundred non-blood-fed (two weeks old) female mosquitoes deprived of water and sugar for at least one hour. Responses were recorded for 5 min; the filter paper rings were swapped or renewed to start another trial. DEET 1% was used as a positive control. Behavioral responses were expressed in percent protection, P%=(C-T)/C)X100 [30, 31].

### Oviposition assays

Three trials were performed. In the first trial, 6 concentrations of fenchone were tested. One hundred μl from each of the primary concentrations of 0.0001, 0.001,0.01,0.1,1, and 10% of fenchone were added to 6 white plastic cups (old-fashioned tumblers, Finelinesettings.com) filled with 100 ml of tap water which would represent in effect a further final 1000x dilution of each primary concentration in each cup. Before adding water and fenchone into each cup, the inside walls were lined with two slightly overlapping 5.8 × 11.4 cm strips of seed germination paper (Regular weight brown seed germination or toweling paper, Seedburo Equipment Company, Des Plaines, IL, USA). The seed germination paper was held against the sides with forceps while pouring the tap water into them to ensure the seed germination paper remained hugged against the cup walls. Control cups contained 100 μl of 100% ethanol added to 100 ml of tap water. Two cups (one with treatment and the other the control cup) were placed in the opposite corners of each of the 6 cages (30.5 cm x 30.5 cm x 30.5 cm; Cat # 1450B Bioquip Products Inc, Rancho Dominguez, CA, USA; discontinued). Eighteen to 21 gravid *Ae. aegypti* females were released into each of the cages. The cages with the mosquitoes and treatment and control cups in them were spaced 128 cm apart in the insectary (80% RH with lighting set at 11 h daylight, 1 h crepuscular, and 12 h dark) for 7 days. Females had constant access to 10% sucrose solution in each cage. On each of the 7 days the dead mosquitoes were counted and removed, and the minimum and maximum temperatures were recorded. The light/dark conditions we simulated in our insectary are conducive for maximum egg-laying activity in *Ae. aegypti* [32]. The concentration of the fenchone remained consistent in each cup for the 7 days as very little evaporation was noted

The second trial with 6 replicates was set up like the first one, except for the tested doses. The control and treatment cups were placed in opposite corners of the cage.

The third trial was set up the same as the first trial but used 24 cages, each with 18-25 gravid females, to test a single dose.

### Field tests

The entire inside walls of black plastic cups (16-oz, 473 ml black stadium cups, csbdstore.com) Charlotte, NC, USA) were lined with rough brown paper (Regular weight seed germination or toweling paper, Seedburo Equipment Company). In each of the six properties in Reedley (Fresno County, CA), traps were placed in shady areas 30 m apart from each other. The control cups were filled with 2.5 ml of ethanol in 247.5 ml of tap water; the infusion cups were prepared with 250 ml 7-day-old Orchard grass infusion; and fenchone cups had 2.5 ml of 1% (+/-)- fenchone solution in ethanol plus 247.5 ml tap water, Seven-day-old infusions were prepared by adding 30 g of Orchard grass to 11.3 L of tap water and left outdoors to ferment in a plastic drum while mixing once a day. The cups were spaced 30 m apart to allow a single female oviposition choice in a corner area of a property. Orchard grass infusion in black cups lined with rough germination paper is used by the local mosquito abatement district for *Ae. aegypti* abundance surveillance purposes. After 7 days, the oviposition paper was removed carefully, and the number of eggs attached to the oviposition paper was counted. This was repeated three times, comprising a field evaluation of 3 × 6 replicates. In each of the three replicates, the cups containing different solutions were placed in alternate positions to avoid possible position effects. In Reedley, *Ae. aegypti* is the only *Aedes* mosquito species that lay eggs in these cups.

The field evaluation was conducted in late summer when the daily average maximum and minimum temperatures were 37 and 19°C. After 7 days when the cups were removed, there were still about 50 ml of infusion or water remaining in all the cups, which meant for the entire 7 days females were able to lay eggs on moist germination paper above the water or infusion line. Cups were left for 7 days in the field because that is the procedure the mosquito abatement district follow for *Ae. aegypti* egg surveillance purposes.

## Acknowledgments

We thank Dr. Pingxi Xu for sharing samples and providing constructive discussions. Research reported in this publication was supported by the National Institute of Allergy and Infectious Diseases (NIAID) of the National Institutes of Health (NIH) under award number R01AI095514. The content is solely the responsibility of the authors and does not necessarily represent the official views of the NIH. Katherine Brisco and AJC acknowledge funding support from the Pacific Southwest Regional Center of Excellence for Vector-Borne Diseases funded by the U.S. Centers for Disease Control and Prevention (Cooperative Agreement 1U01CK000516).

## Notes

### Competing Interest Statement

The authors have declared no competing interest.

